# Area per player in small-sided games to replicate the external load and estimated physiological match demands in elite soccer players

**DOI:** 10.1101/2020.02.03.932954

**Authors:** Andrea Riboli, Giuseppe Coratella, Susanna Rampichini, Emiliano Cé, Fabio Esposito

## Abstract

The current study determined the area-per-player during small- or large-sided games with or without goalkeeper that replicates the relative (m·min^−1^) total distance, high-intensity running distance, sprint distance and metabolic power covered during official matches. Time-motion analysis was performed on twenty-five élite soccer-players during 26 home-matches. A total of 2565 individual samples for SSGs using different pitch sizes and different number of players were collected and classified as SSGs with (SSG-G) or without goalkeeper (SSG-P). A between-position comparison was also performed. The area-per-player needed to replicate the official match demands was *largely* greater in SSG-G *vs* SSG-P for total distance [187±53 vs 115±35 m^2^, effect size (ES): 1.60 95%CI 0.94/2.21], high-intensity running distance [262±72 vs 166±39 m^2^, ES: 1.66(0.99/2.27)] and metabolic power [177±42 vs 94±40, ES: 1.99(1.31/2.67)], but similar for sprint distance [(316±75 vs 295±99 m^2^, ES: 0.24(−0.32/0.79)] with direction of larger area-per-player for sprint distance > high-intensity running > total distance ≅ metabolic power for both SSG-G and SSG-P. In SSG-G, forwards required greater area-per-player than central-defenders [ES: 2.96(1.07/4.35)], wide-midfielders [ES: 2.45(0.64/3.78)] and wide-defenders [ES: 3.45(1.13/4.99)]. Central-midfielders required greater area-per-player than central-defenders [ES: 1.69(0.20/2.90)] and wide-midfielders [ES: 1.35(−0.13/2.57)]. In SSG-P, central defenders need smaller area-per-player (ES: −6.01/−0.92) to overall replicate the match demands compared to all other positions. The current results highlight that soccer players need a specific area-per-player during the small-side games with or without goalkeeper to replicate the overall match demands, especially to perform high-intensity running or sprint distance. Additionally, central defenders, central midfielders and forwards need to be trained with tailored area-per-player or specific rules/additional exercises.

## Introduction

Small- or large-sided games are frequently used to replicate the soccer-specific match demands in terms of technical proficiency, tactical awareness, speed, acceleration/deceleration, and endurance performance [1]. To track these demands, recent tracking technologies such as global positioning systems (GPS) or semi-automatic video-based multi-camera image systems (MCIS), are usually used [2]. In small-sided games (SSGs), the manipulation of pitch size, number of players per team, goalkeeper presence and technical rules modulate the soccer-specific demands depending on the aims of each practice session [1, 3]. Increments in pitch size or reduction in the number of players increases total distance (TD) covered, total high-intensity running distance (HIRD) and total sprint distance (TSD) [4, 5]. Conversely, when pitch size is reduced or the number of players is increased, players get more ball touches but they have not the space to reach the high-speed running, and the total distance covered is rather characterized by acceleration and deceleration (often defined as mechanical work, MechW) [6]. To possibly combine the pitch size and number of players, the area per player (ApP, expressed as m^2^ · player^−1^) has been introduced [1]. Lastly, SSGs can be performed with (SSG-G) or without goalkeepers (SSG-P), when the aim is to out-score the opponent team or to maintain ball possession as long as possible, respectively [1]. Although TD, HIRD and TSD were reviewed to be greater in SSG-P than SSG-G using the same pitch size [7] a subsequent study found greater TSD in SSG-G compared to SSG-P [6] so further investigations are needed [8].

Over the last years, the metabolic power approach has been proposed as a tool to estimate the energetic demands of variable-speed locomotion typically seen in team sports [9, 10]. While it is difficult to measure directly the exact energy cost of changing speed, a metabolic power calculation based on a theoretical model has been used to estimate the energy cost of high-intensity running activities in team sports [9, 11]. However, this model may underestimate very largely the actual net energy demand of soccer-specific exercises, since it does not take into account the actual individual running economy [12, 13]. Notwithstanding, the metabolic power approach could capture the high-demanding locomotor activities independently of the actual speed registered by GPS [13, 14]. Therefore, the combination of the metabolic power and GPS-speed-threshold traditional metrics should be used to present a description of the running demands that include both high speed running and/or accelerations and decelerations [13, 15].

An accurate comparison of the match *vs* training locomotor activities may help to plan the training sessions to replicate the official match demands and to optimize performance goals [5, 16]. Particularly, the locomotor activities during different SSGs compared to official matches are still under investigation. Additionally, discriminating such locomotor activities by position could help to tailor the training session. Therefore, the present study aimed to: i) determine the ApP that could be used to replicate the official matches TD, HIRD, TSD, MechW (normalized as meters covered in one minute) and *P*_met_ (normalized as W·kg^−1^) during both SSG-P and SSG-G; and ii) differentiate the ApP according to playing position. To increase the ecological validity, this was assessed in élite “Serie A” soccer players.

## Materials and Methods

### Subjects

Twenty-five élite soccer players competing in Italian Serie A were involved in the present study (age: 27 ± 5 yrs; body mass: 79 ± 7 kg; body height: 1.84 ± 0.06 m). All participants were classified according to their position: central-defenders (n = 6), wide-defenders (n = 4), central-midfielders (n = 5), wide-midfielders (n = 5) and forwards (n = 5). The goalkeepers were excluded from the data collection. The club’s medical staff certified the health status of each player. An injured player was excluded from the data collection for at least one month after a full in-group re-integration. All participants gave their written consent after a full explanation of the purpose of the study and the experimental design. The local University Ethics Committee approved the study and was performed in accordance with the principles of the Declaration of Helsinki (1975).

### Design

The present investigation was carried out during in-season period (August 2014 – May 2016). The participants undertook their traditional weekly training routine. All sessions were performed on two grass pitches preserved by qualified operators and performed at the same time to limit the effects of circadian variations. A specialized and high-qualified physician staff recommended and monitored the diet regime of each player before and after every training session.

Two different formats of SSGs were analyzed: SSG-G and SSG-P. A total of 2565 (1033 and 1532, respectively) individual GPS samples with a median of 37 (range = 12 to 62) and 56 (range = 25 to 86) in SSG-G and SSG-P respectively were undertaken for each player. The number of players ranged from 5*vs*5 to 10*vs*10, with a pitch area ranging from 800 m^2^ to 6825 m^2^ for SSG-G and 3*v*3 to 10*vs*10 with a pitch area from 400 m^2^ to 4550 m^2^ for SSG-P. Hence, ApP ranged from 67 m^2^ to 341 m^2^ for SSG-G and from 43 m^2^ to 341 m^2^ for SSG-P (for a detailed description of these parameters, see S1 and S2 Tables). ApP was calculated excluding the goalkeepers in SSG-G. Both small- or large-sided games was abbreviated as SSGs and specified by ApP. The SSGs were performed under the supervision and motivation of several coaches to keep up a high work-rate [3]. For the same reason, a ball was always available by prompt replacement when it went out-of-play [1]. In SSG-G, the corners were replaced by a prompt ball-in-game from the goalkeeper [17]. The SSGs were completed after a standardized 20-min warm-up under the club’s staff guidance. Only official home matches (N = 26; individual samples = 228; individual sample range = 6 to 24) were assessed to ensure data consistency [8]. The home-match pitch size was 105 × 66 m, with a grass surface.

To determine the ApP in both SSG-G and SSG-P that replicates the normalized TD, HIRD, TSD, MechW (m·min^−1^) and *P*_met_ (W·kg^−1^) recorded during the official matches, we first recorded these variables during the official matches. Thereafter, we separately plotted each relationship between ApP and the normalized TD, HIRD, TSD, MechW and *P*_met_ during SSG-G or SSG-P. Then, the mean values recorded during the official matches were used to intersect each ApP/ TD, HIRD, TSD, MechW or *P*_met_ relationship recorded in SSG-G or SSG-P to calculate the ApP that corresponded to the official match demands (See Fig 1)

**Fig 1.**
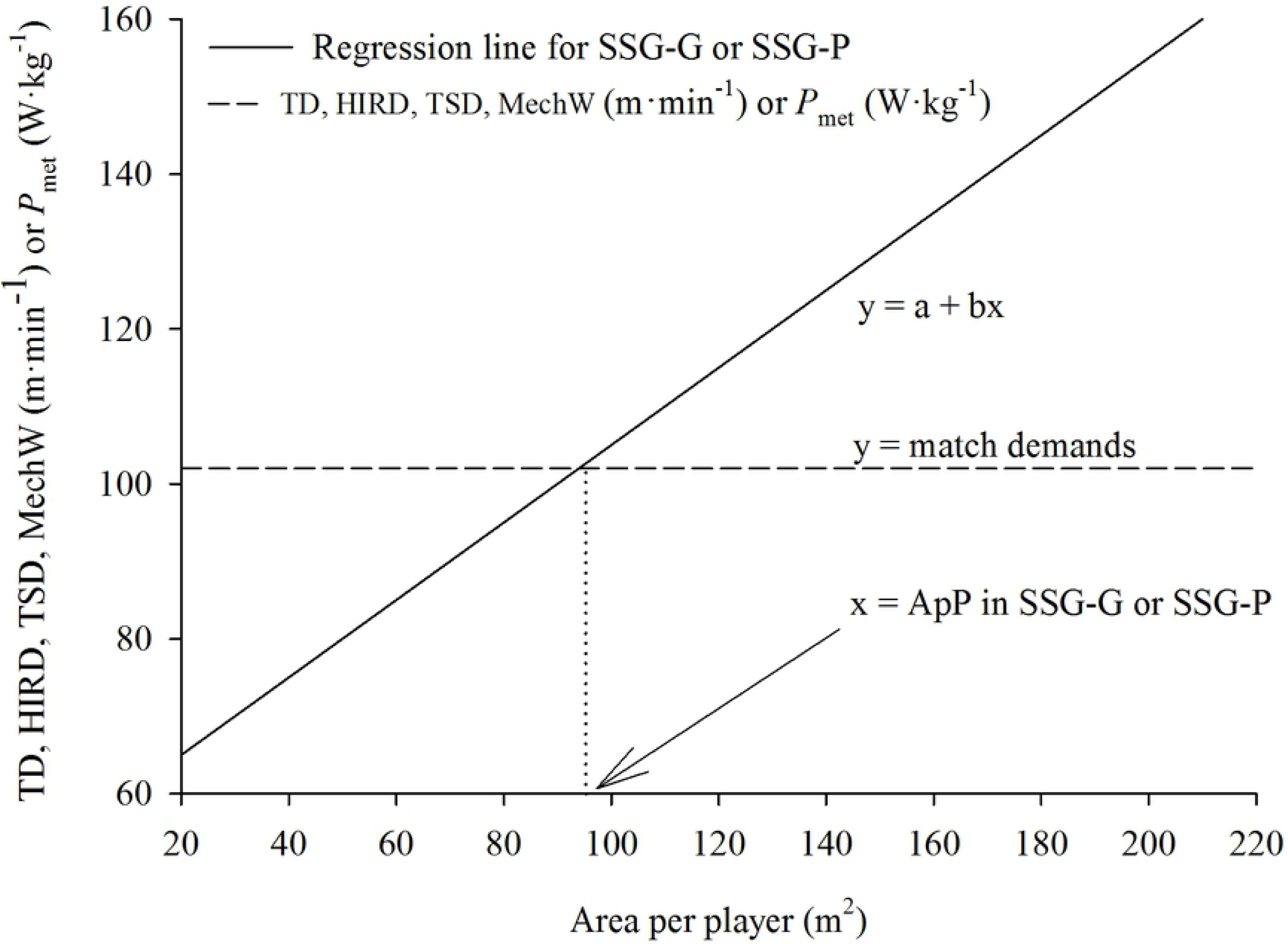
Graphical representation of the procedures used to determine the area per player in SSG-G or SSG-P that matches the official match demands. The area per player in SSG-G or SSG-P is reported on X-axis. The SSG-G or SSG-P demands are reported on Y-axis. The regression line shows how the area per player influences the SSGs demands. The horizontal dashed line represents the official match demands. From the intersection point of the regression line with the horizontal line (i.e. when the SSGs demands equate the official match demands), a vertical dotted line is drawn to the X-axis. The intersection point between the X-axis and the vertical dotted line is the calculated area per player in SSGs necessary to replicate the official match demands.

### Procedures

For the aims of this study, the interchangeability of GPS and MCIS for TD, HIRD, TSD, MechW and *P*_met_ needed to be calculated as first step. A 10Hz GPS (K-Sport, Montelabbate, Italy) unit was used to collect data during the training sessions [18]. The GPS unit was placed within a dedicated pouch between the player’s shoulder blades (upper thoracic-spine) in a sports vest and worn under the playing jersey. Each device was turned on at least 15-min before each data collection to allow the acquisition of the satellite signal [6]. To reduce the inter-unit differences, every player wore the same device for each training session over the whole investigation [19]. The locomotor activities during the official matches were collected using a computerized semi-automated MCIS (Prozone Sport, Leeds, UK) and processed by a dedicated software (K-SportOnline, K-Sport, Montelabbate, Italy). The system has previously been shown to provide valid and reliable measurements of the match activity in soccer [20, 21].

During both training sessions and home-matches, total distance, total high-intensity running distance (>15 km·h^−1^), total sprint distance (>24 km·h^−1^) [3, 8, 21]. Additionally, total mechanical work distance (MechW,) i.e. the total distance of velocity changes calculated using >2 m·s^−2^ accelerations and decelerations was calculated [4, 5]. The average metabolic power (*P*_met_) was calculated following previous procedures [10, 22]. TD, HIRD, TSD and MechW were normalized as relative distance covered in one minute (m·min^−1^), while *P*_met_ were normalized as watt per kilogram (W·kg^−1^); than all parameters were inserted into the data analysis.

TD, HIRD, TSD and MechW were measured using either GPS or the MCIS. Therefore, to check the interchangeability of these two tracking technologies, a simulated match was monitored using both systems simultaneously [2]. All data were collected in the stadium where the official matches were played and locomotor data were integrated with calibration equations as proposed by other authors [2].

### Statistical analysis

Statistical analysis was performed using a statistical software package (SigmaPlot v-12.5, Systat Software Inc., San Jose, CA, USA). To check the normal distribution of the sampling, a Shapiro-Wilk test was used. The Pearson’s product moment and the typical error of the estimate (TEE) were calculated to determine the relationship between the GPS and the MCIS. The correlation coefficient was interpreted as follows: *r* =0.00-0.09 *trivial*, 0.10-0.29 *small*, 0.30-0.49 *moderate*, 0.50-0.69 *large*, 0.70-0.89 *very large*, 0.90-0.99 *nearly perfect*; the threshold values for the TEE were interpreted as follows: >0.2 *small*, >0.6 *moderate*, >1.2 *large* and >2 *very large* [23]. A linear regression analysis was used to calculate the correlation between TD, HIRD, TSD, MechW, *P*_met_ and the ApP during both SSG-G and SSG-P. Thereafter, a two-way ANOVA was used to calculate the difference in the optimal ApP in TD, HIRD, TSD, MechW, *P*_met_ calculated for SSG (SSG-G *vs* SSG-P) and position (central-defenders, wide-defenders, central-midfielders, wide-midfielders and forwards). A post-hoc analysis (Holm-Sidak correction) was used to calculate the differences in the independent factors. The effect size with 95% confidence intervals (CI) was calculated and interpreted as follows: <0.20: *trivial*; 0.20-0.59: *small;* 0.60-1.19: *moderate;* 1.20-1.99: *large*; ≥2.00: *very large* [23]. Statistical significance was set at α < 0.05. Unless otherwise stated, all values are presented as mean ± standard deviation (SD).

## Results

The magnitude of the GPS *vs* MCIS bias were *trivial* for TD (−3.0 ± 1.3%, ES = −0.18, CI: – 0.80/0.44), HIRD (−3.3 ± 1.6%, ES = −0.12, CI: −0.74/0.51), TSD (−3.9 ± 10.9%, ES = −0.11, CI: −0.44/0.22) and MechW (−4.1 ± 17.9%, ES = −0.19, CI: −0.80/0.44) and small for *P*_met_ (−4.0 ± 0.6%, ES = −0.38, CI: −1.27/0.49). A *small* TEE was found between the MCIS and GPS for TD (TEE: 0.09, CI: 0.07/0.14), HIRD (TEE: 0.04, CI: 0.03/0.06), TSD (TEE: 0.08, CI: 0.07/0.10) and *P*_met_ (TEE: 0.07, CI: 0.04/0.13), while a *moderate* TEE was found for MechW (TEE: 0.75, CI: 0.56/1.10). In addition, a *nearly perfect* correlation was observed for TD, HIRD, TSD and *P*_met_ (r =0.99, r =0.98, r =0.99 and r = 0.98 respectively; p <0.001), and a *moderate* correlation for MechW (r =0.67; P =0.002) measured using GPS and MCIS.

As shown in Figs 2 and 3, in SSG-P a *very large* correlation between the relative distance and the ApP was found for TD, HIRD, TSD and *P*_met_ while a *moderate* correlation was found for MechW. In SSG-G, a *very large* correlation between the relative distance and the ApP for TD, HIRD and TSD, a *large* correlation for *P*_met_ and a *moderate* negative correlation for MechW were found. Because of the *moderate* correlations observed for MechW in both SSG-G and SSG-P, we did not perform the calculation or the ApP for MechW, given the high risk of bias.

**Fig 2.**
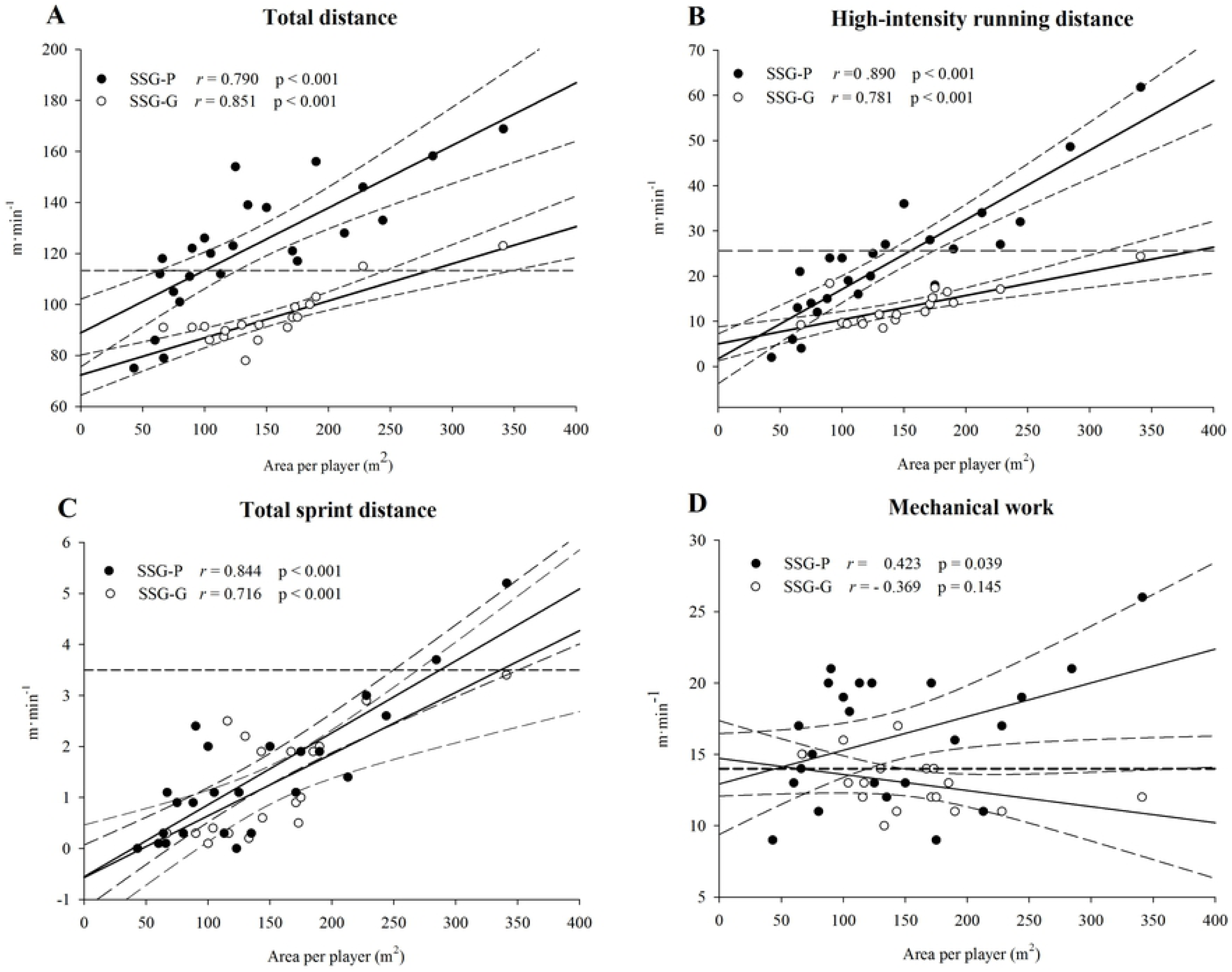
Relationship between area per player and relative speed distances. The relationship between the area per player (m^2^·player) and relative distance (m·min^−1^) during both small-sided games possession-play without goalkeepers (SSG-P, closed circles) and small-sided games with goalkeepers (SSG-G, open circles) is shown for total distance (**A**), high-intensity running distance (**B**), total sprint distance (**C**) and mechanical work (**D**). A linear regression analysis with 95% confidence bands is also reported for SSG-P and SSG-G. The horizontal dashed line represents the average value measured during the official matches. The correlation between the area per player and the relative distance for both SSG-P and SSG-G is also reported.

**Fig 3.**
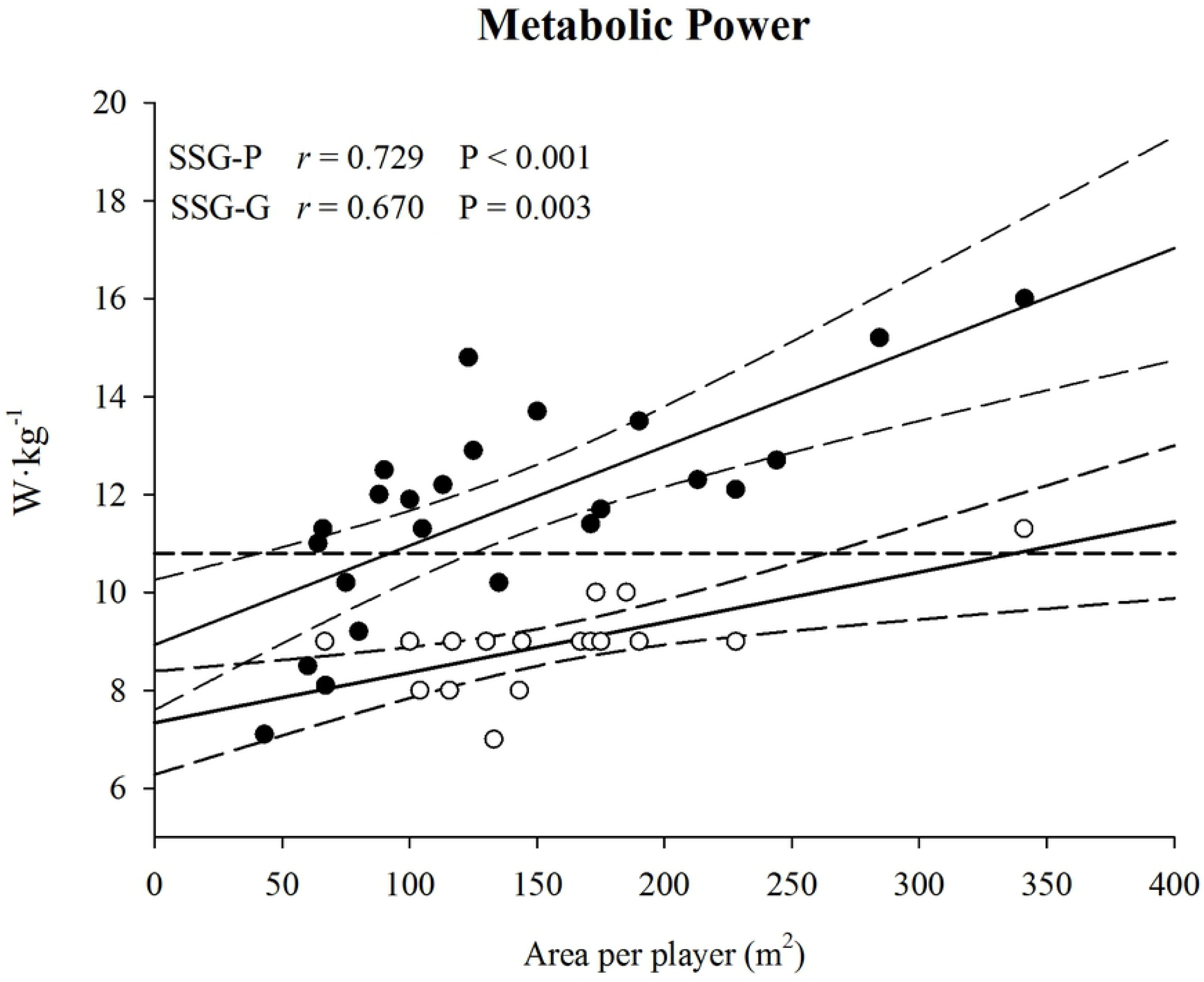
Relationship between area per player and estimated metabolic power. The relationship between the area per player (m^2^·player) and average metabolic power (W·kg^−1^) during both small-sided games possession-play without goalkeepers (SSG-P, closed circles) and small-sided games with goalkeepers (SSG-G, open circles) is shown. A linear regression analysis with 95% confidence bands is also reported for SSG-P and SSG-G. The horizontal dashed line represents the average value measured during the official matches. The correlation between the area per player and the relative distance for both SSG-P and SSG-G is also reported.

For both SSG-P and SSG-G, the ApP necessary to replicate the relative distance recorded during the matches for TD, HIRD, TSD and *P*_met_ is shown in table 1. No SSG × position interaction was found (p =0.674) for ApP for TD. A main effect for SSG (p <0.001) and position (p =0.024) was detected. The between-SSG post-hoc analysis is reported in table 1. In SSG-P, a larger ApP is required for forwards *vs* central-defenders (p =0.023; ES = 4.35, CI: 1.93/6.01), with no other between-position differences. In SSG-G, no between-position difference occurred.

**Table 1.** Area per player to replicate official-match load using SSGs. The area per player (m^2^·player) to replicate the same amount of total distance (TD, m·min ^−1^), high intensity running distance (HIRD, m·min^−1^), sprint distance (TSD, m·min^−1^) and average metabolic power (*P*_met_, W·kg^−1^) as recorded in the matches is shown for the team average (Total) and split for central defenders (CD, n = 6), wide defenders (WD, n = 4), central midfielders (CM, n = 5), wide midfielders (WM, n = 5) and forwards (FW, n = 5). The area per player is reported for both SSG-P and SSG-G. Data are presented as mean(SD); P values, effect size (ES) and confidence intervals (CI) are reported.

| | TD | | | | HIRD | | | | TSD | | | | $P_{\text{met}}$ | | | |
| --- | --- | --- | --- | --- | --- | --- | --- | --- | --- | --- | --- | --- | --- | --- | --- | --- |
| Position | SSG-P | SSG-G | p | ES (CI) | SSG-P | SSG-G | p | ES (CI) | SSG-P | SSG-G | p | ES (CI) | SSG-P | SSG-G | p | ES (CI) |
| Total | 115(35) | 187(53)* | <0.001 | -1.60 (-2.21/-0.94) | 166(39) | 262(72)* | <0.001 | -1.66 (-2.27/-0.99) | 295(99) | 316(75) | 0.415 | -0.24 (-0.79/0.32) | 94(40) | 177(42)* | <0.001 | -1.99 (-2.67/-1.31) |
| CD | 65(24)<sup>#</sup> | 165(26)* | <0.001 | -4.00 (-5.55/-1.83) | 122(30)<sup>#</sup> | 205(57)*<sup>#§</sup> | <0.001 | -1.82 (-3.00/-0.37) | 257(76) | 278(51) | 0.672 | -0.32 (-1.44/0.84) | 31(11) | 151(23)* | <0.001 | -6.14 (-8.85/-3.44) |
| WD | 121(21) | 193(71)* | 0.023 | -1.38 (-2.70/0.31) | 163(30) | 246(36)*<sup>#</sup> | 0.003 | -2.50 (-3.92/-0.43) | 297(26) | 274(67) | 0.696 | 0.45 (-1.01/1.79) | 106(31)<sup>°</sup> | 183(27)* | 0.003 | -2.39 (-4.01/-0.77) |
| CM | 119(9) | 184(41)* | 0.021 | -2.12 (-3.41/-0.42) | 174(28) | 311(69)* | <0.001 | -2.60 (-3.96/-0.74) | 329(66) | 340(33) | 0.834 | -0.21 (-1.43/1.05) | 107(13)<sup>°</sup> | 191(25)* | <0.001 | -3.81 (-5.88/-1.73) |
| WM | 135(20) | 183(81)* | 0.079 | -0.81 (-2.02/0.55) | 172(19) | 222(72)*<sup>#§</sup> | 0.031 | -0.95 (-2.15/0.44) | 264(52) | 281(62) | 0.758 | -0.30 (-1.51/0.98) | 132(12)<sup>°</sup> | 180(61)* | 0.047 | -0.95 (-2.71/0.51) |
| FW | 147(9) | 214(51)* | 0.018 | -1.83 (-3.09/-0.22) | 207(28) | 333(12)* | <0.001 | -5.85 (-7.91/-2.66) | 334(92) | 407(68) | 0.176 | -0.90 (-2.10/0.48) | 99(2)<sup>°</sup> | 201(66)* | <0.001 | -1.97 (-3.48/-0.46) |

\*: $p < 0.05$ vs SSG-P, #: $p < 0.05$ vs forwards, §: $p < 0.05$ vs central midfielders, ° $p < 0.05$ vs central defenders

No SSG × position interaction was found (p =0.065) for ApP for HIRD. A main effect for SSG (p <0.001) and position (p <0.001) was detected. The between-exercise post-hoc analysis is reported in table 1. In SSG-P, a greater ApP is required for forwards *vs* central-defenders (p =0.024; ES = 2.92, CI: 1.04/4.29), with no other between-position differences. In SSG-G, forwards required greater ApP than central-defenders (p <0.001; ES = 2.96, CI: 1.07/4.35), wide-midfielders (p =0.002; ES = 2.45, CI: 0.64/3.78) and wide-defenders (p =0.029, ES = 3.45, CI: 1.13/4.99). Central-midfielders required a greater ApP than central-defenders (p =0.002; ES = 1.69, CI: 0.20/2.90) and wide-midfielders (p =0.019, ES = 1.35, CI: 0.13/2.57).

No SSG × position interaction was found (p =0.803) for ApP for TSD, not even a main effect for exercise (p =0.415). A main effect for position (p =0.049) was detected. The between-exercise post-hoc analysis is reported in table 1. In both SSG-P and SSG-G, no between-position difference occurred.

No SSG × position interaction was found (p =0.167) for ApP for *P*_met_. A main effect for SSG (p <0.001) and position (p =0.002) was detected. The between-SSG post-hoc analysis is reported in table 1. In SSG-P, a smaller ApP is required for central-defenders *vs* wide-defenders (p =0.031; ES = −2.69, CI: −4.32/−1.05), wide-midfielders (p <0.001; ES = −2.64, CI: −4.35/−0.92), central-midfielders (p =0.028; ES = −5.10, CI: −7.53/−2.66), forwards (p =0.024; ES = – 1.89, CI: −3.32/0.47). In SSG-G, no between-position difference occurred.

## Discussion

### Methodological considerations

Some preliminary considerations need to be done to properly interpret the results observed here. The interchangeability between the GPS and MCIS needs to be carefully checked, especially when recording high-speed or non-linear movements [2]. Notwithstanding, the present results are based on the *trivial* differences in the metrics recorded using either the GPS or MCIS. Additionally, the current Serie-A stadia’ structures could not allow accurate GPS acquisition.

Secondly, The *P*_met_ represents an estimation of the energy expenditure based on variations in speed, i.e., acceleration and deceleration [10]. The *P*_met_ depends on the amount of the GPS metrics, and may be used to provide an overall description of the energy demands. However, *P*_met_ may underestimate very largely the actual net energy demands if compared with traditional calorimetry assessment [13]. However, due to technological limitation during training sessions/matches, *P*_met_ is the only way to estimate with satisfactory accuracy the match demands [9]. Additionally, when *P*_met_ is combined with the GPS speed-threshold traditional metrics (TD, HIRD and TSD), this provides more detailed information about the actual match or training mechanical load (i.e., high-speed *vs.* acceleration/deceleration) [13].

### Main findings

The first novel finding observed in the present study was a detailed calculation of the ApP in SSG-P or SSG-G necessary to replicate the TD, HIRD, TSD and *P*_met_ recorded during the official matches. It is here shown that, irrespective of the SSG type, the higher the speed threshold, the larger the ApP required (i.e., TSD > HIRD > TD ≈ *P*_met_). Secondly, the inclusion of the goalkeeper increases the ApP for TD, HIRD and *P*_met_, while no difference was observed in SSG-P *vs* SSG-G for TSD. Additionally, central defenders resulted in the smallest ApP compared to all other positions, both in SSG-P and SSG-G. Lastly, both central-midfielders and forwards need the largest ApP compared to all other positions, both in SSG-G and SSG-P, to replicate the match demands.

In combination with high-intensity physiologically based exercises that help soccer players adapt their physiology to the demands of playing an official match [24], determining the ApP to be used during the training sessions that replicates the match external-load demands may help sport physiologists and practitioners to plan properly the weekly training load for specific performance goal [5]. Both in SSG-P and in SSG-G, larger ApP leads to greater distance covered whatever the speed threshold [5]. Accordingly, the present findings highlight that the ApP in SSGs to replicate the TSD match demands is very close to the official match ApP (≈ 340 m^2^). In line with the present outcomes, it was shown that the larger the pitch size, the greater the distance covered at speed >18 km·h^−1^ [25]. Other authors found that TD and the distance covered at 19.8–25.2 km·h^−1^ and >25.2 km·h^−1^ increased proportionally with the pitch size [26]. A recent study reported that ApP ∼ 311 m^2^ was able to replicate the high-speed match demands during SSG-G [5]. The exposure to high-demanding activities was shown to improve the players’ fitness level, to prepare the players to the match workload and to result in a greater protection against non-contact injuries [27-29]. Therefore, manipulating properly the ApP allows managing the training loads to be performed using SSGs, both for performance and prevention purposes.

The current outcomes also highlight that training using SSGs with or without goalkeeper affects the ApP necessary to replicate the official match demands. Particularly, with the exception of TSD, the goalkeeper presence increases the ApP for TD, HIRD and *P*_met_, i.e. SSG-G > SSG-P. Partially in contrast with the present outcomes, it was reported that SSG-G resulted in greater TSD than found in SSG-P [6]. However, the authors investigated a maximum ApP of 135 m^2^, hence, this does not allow an appropriate comparison. Also other researchers reported that the TD and the time spent in high-intensity running (>17 km·h^−1^) was greater with goalkeepers [30]. Although the authors argued that the goalkeeper presence might have motivated the players, two subsequent reviews [1, 7] consistently remarked that the goalkeeper presence could improve the players’ organization, thus decreasing the SSGs demands. Indeed, during SSG-G, the two teams’ aim is to outscore the opponent team, while maintaining a match-like tactical organization. In contrast, since during SSG-P the aim is to maintain the ball possession as long as possible, the players are freer to move across the selected pitch size. This rule-difference seems to account for the largest ApP in SSG-G necessary to replicate the TD, HIRD and *P*_met_ recorded during the official matches.

To our knowledge, the calculation of the ApP across positions was used here for the first time. No between-position difference in the ApP was found for TSD, neither in SSG-P nor SSG-G. In SSG-P, it was observed that central defenders need smaller ApP than forwards for TD, and HIRD, while smaller ApP than all other position for *P*_met_. In SSG-G, no between-position difference in ApP was observed for TD, TSD and *P*_met_, while forwards and central midfielders need larger ApP than central defenders and wide midfielders for HIRD suggesting that these positions should be conditioned differently during SSGs. Defenders used to move within a “defined” space over the official match, while central-midfielders and forwards move across the pitch to gain possession of the ball, marking the opponent or creating space to score [21]. This might be considered for the smaller ApP needed to accumulate the match demands in central defenders than forwards/central-midfielders. However, the sprinting activities are not influenced by position, since these appear to need large pitch areas available anyhow. Thus, some positions need larger ApP or specific rules/additional exercises to replicate the HIRD or TSD accumulated over the matches.

The present outcomes could be used in practice to: i) calculate an ApP that replicate an estimated match demand using *P*_met_ for both SSG-P and SSG-G; ii) replicate the official relative match demands using the specific minimal ApP to HIRD or TSD be accumulated during the SSG-G/P performed in the training sessions; iii) differentiate the ApP when SSG-P or SSG-G are performed according to the aim of the training session; iv) differentiate the ApP between position when needed or propose specific additional exercises or rules to overload or unload each player.

## Conclusions

In conclusion, the present novel findings were the first to investigate the ApP during the SSGs required to elicit the official match demands. The current results highlight that soccer players need a specific area-per-player during the SSGs with or without goalkeeper to replicate the overall match demands, especially to perform HIRD or TSD. Additionally, central defenders, central midfielders and forwards need to be trained with tailored area-per-player or specific rules/additional exercises. Taken together with more physiologically based exercises, these results allow to manage the training loads towards the desired players’ fitness component to maximize transfer to the game-like and performance goal using SSGs.

## Acknowledgments

The authors report no conflict of interest.

## Supporting information

**S1 Table. Small-sided games with goalkeepers (SSG-G).** The SSG-G recorded during the 2-season training sessions are shown. The SSG-G are split for the number of players and pitch size (width × length). The total pitch area (m^2^) and area per player (ApP, m^2^·player) have been calculated. The average number of observations per player (mean, max-min) for each condition are also reported as mean(SD).

**S2 Table. Small-sided games without goalkeepers (SSG-P).** The SSG-P recorded during the 2-season training sessions are shown. The SSG-P are split for the number of players and pitch size (width × length). The total pitch area (m^2^) and area per player (ApP, m^2^·player) have been calculated. The average number of observations per player (mean, max-min) for each condition are also reported as mean(SD).

## References

1. Hill-Haas SV, Dawson B, Impellizzeri FM, Coutts AJ. Physiology of small-sided games training in football: a systematic review. Sports Med. 2011;41(3):199–220. Epub 2011/03/15. doi: 10.2165/11539740-000000000-00000. PubMed PMID: 21395363.

2. Buchheit M, Allen A, Poon TK, Modonutti M, Gregson W, Di Salvo V. Integrating different tracking systems in football: multiple camera semi-automatic system, local position measurement and GPS technologies. J Sports Sci. 2014;32(20):1844–57. Epub 2014/08/06. doi: 10.1080/02640414.2014.942687. PubMed PMID: 25093242.

3. Rampinini E, Coutts AJ, Castagna C, Sassi R, Impellizzeri FM. Variation in top level soccer match performance. Int J Sports Med. 2007;28(12):1018–24. Epub 2007/05/15. doi: 10.1055/s-2007-965158. PubMed PMID: 17497575.

4. Lacome M, Simpson BM, Cholley Y, Buchheit M. Locomotor and Heart Rate Responses of Floaters During Small-Sided Games in Elite Soccer Players: Effect of Pitch Size and Inclusion of Goalkeepers. Int J Sports Physiol Perform. 2018;13(5):668–71. Epub 2017/09/28. doi: 10.1123/ijspp.2017-0340. PubMed PMID: 28952828.

5. Lacome M, Simpson BM, Cholley Y, Lambert P, Buchheit M. Small-Sided Games in Elite Soccer: Does One Size Fit All? Int J Sports Physiol Perform. 2018;13(5):568–76. Epub 2017/07/18. doi: 10.1123/ijspp.2017-0214. PubMed PMID: 28714774.

6. Gaudino P, Alberti G, Iaia FM. Estimated metabolic and mechanical demands during different small-sided games in elite soccer players. Hum Mov Sci. 2014;36:123–33. Epub 2014/06/27. doi: 10.1016/j.humov.2014.05.006. PubMed PMID: 24968370.

7. Aguiar M, Botelho G, Lago C, Macas V, Sampaio J. A review on the effects of soccer small-sided games. J Hum Kinet. 2012;33:103–13. Epub 2013/03/15. doi: 10.2478/v10078-012-0049-x. PubMed PMID: 23486554; PubMed Central PMCID: PMCPMC3588672.

8. Gregson W, Drust B, Atkinson G, Salvo VD. Match-to-match variability of high-speed activities in premier league soccer. Int J Sports Med. 2010;31(4):237–42. Epub 2010/02/17. doi: 10.1055/s-0030-1247546. PubMed PMID: 20157871.

9. Polglaze T, Hoppe MW. Metabolic Power: A Step in the Right Direction for Team Sports. Int J Sports Physiol Perform. 2019:1–5. Epub 2019/02/09. doi: 10.1123/ijspp.2018-0661. PubMed PMID: 30732493.

10. di Prampero PE, Botter A, Osgnach C. The energy cost of sprint running and the role of metabolic power in setting top performances. Eur J Appl Physiol. 2015;115(3):451–69. Epub 2015/01/01. doi: 10.1007/s00421-014-3086-4. PubMed PMID: 25549786.

11. Minetti AE, Pavei G. Update and extension of the ‘equivalent slope’ of speed-changing level locomotion in humans: a computational model for shuttle running. J Exp Biol. 2018;221(Pt 15). Epub 2018/06/14. doi: 10.1242/jeb.182303. PubMed PMID: 29895678.

12. Brown DM, Dwyer DB, Robertson SJ, Gastin PB. Metabolic Power Method: Underestimation of Energy Expenditure in Field-Sport Movements Using a Global Positioning System Tracking System. Int J Sports Physiol Perform. 2016;11(8):1067–73. Epub 2016/03/22. doi: 10.1123/ijspp.2016-0021. PubMed PMID: 26999381.

13. Buchheit M, Manouvrier C, Cassirame J, Morin JB. Monitoring Locomotor Load in Soccer: Is Metabolic Power, Powerful? Int J Sports Med. 2015;36(14):1149–55. Epub 2015/09/24. doi: 10.1055/s-0035-1555927. PubMed PMID: 26393813.

14. Coutts AJ, Kempton T, Sullivan C, Bilsborough J, Cordy J, Rampinini E. Metabolic power and energetic costs of professional Australian Football match-play. J Sci Med Sport. 2015;18(2):219–24. Epub 2014/03/05. doi: 10.1016/j.jsams.2014.02.003. PubMed PMID: 24589369.

15. Stevens TG, De Ruiter CJ, Van Maurik D, Van Lierop CJ, Savelsbergh GJ, Beek PJ. Measured and estimated energy cost of constant and shuttle running in soccer players. Med Sci Sports Exerc. 2015;47(6):1219–24. Epub 2014/09/12. doi: 10.1249/MSS.0000000000000515. PubMed PMID: 25211365.

16. Martin-Garcia A, Gomez Diaz A, Bradley PS, Morera F, Casamichana D. Quantification of a Professional Football Team’s External Load Using a Microcycle Structure. J Strength Cond Res. 2018. Epub 2018/09/11. doi: 10.1519/JSC.0000000000002816. PubMed PMID: 30199452.

17. Castellano J, Casamichana D, Dellal A. Influence of game format and number of players on heart rate responses and physical demands in small-sided soccer games. J Strength Cond Res. 2013;27(5):1295–303. doi: 10.1519/JSC.0b013e318267a5d1. PubMed PMID: 22836601.

18. Castagna C, D’Ottavio S, Cappelli S, Araujo Povoas SC. The Effects of Long Sprint Ability-Oriented Small-Sided Games Using Different Ratios of Players to Pitch Area on Internal and External Load in Soccer Players. Int J Sports Physiol Perform. 2019:1265–72. Epub 2019/03/13. doi: 10.1123/ijspp.2018-0645. PubMed PMID: 30860405.

19. Buchheit M, Al Haddad H, Simpson BM, Palazzi D, Bourdon PC, Di Salvo V, et al. Monitoring accelerations with GPS in football: time to slow down? Int J Sports Physiol Perform. 2014;9(3):442–5. doi: 10.1123/ijspp.2013-0187. PubMed PMID: 23916989.

20. Rampinini E, Alberti G, Fiorenza M, Riggio M, Sassi R, Borges TO, et al. Accuracy of GPS devices for measuring high-intensity running in field-based team sports. Int J Sports Med. 2015;36(1):49–53. doi: 10.1055/s-0034-1385866. PubMed PMID: 25254901.

21. Di Salvo V, Baron R, Tschan H, Calderon Montero FJ, Bachl N, Pigozzi F. Performance characteristics according to playing position in elite soccer. Int J Sports Med. 2007;28(3):222–7. Epub 2006/10/07. doi: 10.1055/s-2006-924294. PubMed PMID: 17024626.

22. Osgnach C, Poser S, Bernardini R, Rinaldo R, di Prampero PE. Energy cost and metabolic power in elite soccer: a new match analysis approach. Med Sci Sports Exerc. 2010;42(1):170–8. Epub 2009/12/17. doi: 10.1249/MSS.0b013e3181ae5cfd. PubMed PMID: 20010116.

23. Hopkins WG, Marshall SW, Batterham AM, Hanin J. Progressive statistics for studies in sports medicine and exercise science. Med Sci Sports Exerc. 2009;41(1):3–13. Epub 2008/12/19. doi: 10.1249/MSS.0b013e31818cb278. PubMed PMID: 19092709.

24. Helgerud J, Engen LC, Wisloff U, Hoff J. Aerobic endurance training improves soccer performance. Med Sci Sports Exerc. 2001;33(11):1925–31. Epub 2001/11/02. doi: 10.1097/00005768-200111000-00019. PubMed PMID: 11689745.

25. Hill-Haas SV, Dawson BT, Coutts AJ, Rowsell GJ. Physiological responses and timemotion characteristics of various small-sided soccer games in youth players. J Sports Sci. 2009;27(1):1–8. Epub 2008/11/08. doi: 10.1080/02640410902761199. PubMed PMID: 18989820.

26. Gaudino P, Iaia FM, Alberti G, Hawkins RD, Strudwick AJ, Gregson W. Systematic bias between running speed and metabolic power data in elite soccer players: influence of drill type. Int J Sports Med. 2014;35(6):489–93. Epub 2013/10/30. doi: 10.1055/s-0033-1355418. PubMed PMID: 24165959.

27. Bowen L, Gross AS, Gimpel M, Li FX. Accumulated workloads and the acute:chronic workload ratio relate to injury risk in elite youth football players. Br J Sports Med. 2017;51(5):452–9. Epub 2016/07/28. doi: 10.1136/bjsports-2015-095820. PubMed PMID: 27450360; PubMed Central PMCID: PMCPMC5460663.

28. Malone S, Roe M, Doran DA, Gabbett TJ, Collins K. High chronic training loads and exposure to bouts of maximal velocity running reduce injury risk in elite Gaelic football. J Sci Med Sport. 2017;20(3):250–4. Epub 2016/08/25. doi: 10.1016/j.jsams.2016.08.005. PubMed PMID: 27554923.

29. Bowen L, Gross AS, Gimpel M, Bruce-Low S, Li FX. Spikes in acute:chronic workload ratio (ACWR) associated with a 5-7 times greater injury rate in English Premier League football players: a comprehensive 3-year study. Br J Sports Med. 2019. Epub 2019/02/23. doi: 10.1136/bjsports-2018-099422. PubMed PMID: 30792258.

30. Dellal A, Chamari K, Pintus A, Girard O, Cotte T, Keller D. Heart rate responses during small-sided games and short intermittent running training in elite soccer players: a comparative study. J Strength Cond Res. 2008;22(5):1449–57. Epub 2008/08/21. doi: 10.1519/JSC.0b013e31817398c6. PubMed PMID: 18714244.

